# Single-Stranded DNA with Internal Base Modifications Mediates Highly Efficient Gene Insertion in Primary Cells

**DOI:** 10.1101/2024.02.01.578476

**Authors:** Karen L. Kanke, Rachael E. Rayner, Eli Abel, Aparna Venugopalan, Ma Suu, Jacob T. Stack, Reza Nouri, Gongbo Guo, Tatyana A. Vetter, Estelle Cormet-Boyaka, Mark E. Hester, Sriram Vaidyanathan

## Abstract

Single-stranded DNA (ssDNA) templates along with Cas9 have been used for gene insertion but suffer from low efficiency. Here, we show that ssDNA with chemical modifications in 10-17% of internal bases (eDNA) is compatible with the homologous recombination machinery. Moreover, eDNA templates improve gene insertion by 2-3 fold compared to unmodified and end-modified ssDNA in airway basal stem cells (ABCs), hematopoietic stem and progenitor cells (HSPCs), T-cells and endothelial cells. Over 50% of alleles showed gene insertion in three clinically relevant loci (*CFTR, HBB*, and *CCR5*) in ABCs using eDNA and up to 70% of alleles showed gene insertion in the *HBB* locus in HSPCs. This level of correction is therapeutically relevant and is comparable to adeno-associated virus-based templates. Knocking out TREX1 nuclease improved gene insertion using unmodified ssDNA but not eDNA suggesting that chemical modifications inhibit TREX1. This approach can be used for therapeutic applications and biological modeling.

## INTRODUCTION

CRISPR/Cas9-based genome editing has been widely exploited for therapeutic purposes and for modeling the biological impact of genetic variants.(1,2) Cas9 nuclease directed to a targeted genomic locus by a single guide RNA (sgRNA) can induce targeted double-stranded breaks (DSBs) that are repaired by DNA repair mechanisms; non-homologous end joining (NHEJ), microhomology-mediated end joining (MMEJ) and homology-directed repair (HR). NHEJ and MMEJ are error-prone pathways that result in insertions and deletions (INDELs) of nucleotides that can knock out (KO) a gene. HR can be used to precisely insert exogenous genetic sequences if a template containing regions homologous to the DSB site is delivered into the cells of interest. Among the different HR template delivery strategies, delivery using adeno-associated viruses (AAV) results in the most efficient gene insertion in multiple cell types including hematopoietic stem and progenitor cells (HSPC), T-cells, airway basal stem cells (ABCs), pluripotent stem cells and mesenchymal cells.(3-8) Cas9 and AAV have been used to make gene edited cell therapies for therapeutic applications.(9) Unfortunately, a recent trial testing the use of hematopoietic stem and progenitor cells (HSPCs) gene edited using Cas9/AAV to treat sickle cell disease was paused because of abnormally low blood cell counts following the transplantation of corrected HSPCs.(10,11) Indeed, previous studies in HSPCs have reported reduced colony formation and engraftment in mouse models that has been attributed to TP53 activation by the AAV genomes.(8,10,12) In addition, integration of inverted terminal repeats (ITRs) from the AAV genome has also been reported in studies investigating in vivo gene editing.(13) Apart from therapeutic applications, gene editing using Cas9/AAV has been used to model the impact of mutations in genetic disorders such as CF and cancer.(14,15) The generation of AAV based templates requires cloning followed by vector production which can take several weeks and thus limit the throughput of such experiments. Thus, adverse outcomes in clinical trials, the observation of aberrant genetic changes and the desire to generate primary cell models quickly have all increased the interest in readily available, alternative HR templates that enable high efficiency gene insertion.

Short ssDNA has been used as a readily available non-viral alternative to AAV-based HR templates. However, the efficiency of gene insertion was significantly lower than AAV-based templates in ABCs and HSPCs in previous studies.(8,16) This is likely due to the premature degradation of ssDNA within the cells by intracellular nucleases. Studies have reported that chemically modifying the ends of ssDNA improves gene correction in cell lines.(17) However, end-modification of ssDNA did not improve gene insertion in hematopoietic stem cells.(18) Cas9 Target Sequences (CTS) that improve the nuclear transport of ssDNA have been reported.(19) However, CTS needs to be designed separately for each sgRNA and creates room for errors in the design process. Thus, a simpler approach to obtain highly efficient non-viral gene insertion without requiring any specialized design or optimization can have many applications.

The modification of internal bases has been shown to improve sgRNA and mRNA activity.(20-23) A similar approach has not been reported for improving HR using ssDNA templates. We reasoned that chemical modification of internal bases may improve the protection of ssDNA from nuclease-mediated degradation. However, it was unclear if the strategy would improve gene insertion because the internal base modifications that inhibit nuclease degradation may also prevent the templates from being recognized by the proteins mediating HR. Furthermore, the approach could be rendered non-viable if the modified bases reduce the fidelity of the HR process and thus introduce missense mutations.

In this study, we tested the use of ssDNA with different proportions of chemically modified bases in ABCs, HSPCs, human vascular endothelial cells (HUVECs), T-cells, and induced pluripotent stem cells (iPSCs). We tested the use of phosphorothioate and 2’-O-methyl modifications that have been used to protect sgRNA and antisense oligonucleotide therapeutics.(24,25) We began our study with a focus on ABCs and attempted to correct mutations associated with cystic fibrosis (CF). CF is caused by mutations in the cystic fibrosis transmembrane conductance regulator (*CFTR*) gene. Although CF is a systemic disorder affecting many organs, the major cause of mortality is lung failure caused by recurrent lung infections. Gene edited airway stem cell therapies to correct the most common CF causing mutation (F508del) have been proposed and Cas9/AAV have been used to correct the F508del mutation with high efficiency.(5,26) We began by evaluating correction of the F508del mutation in ABCs. To test the widespread applicability of this approach, we further extended these experiments to other therapeutically relevant loci (*HBB, CCR5*) in ABCs and other cell types such as HSPCs, T-cells, HUVECs and iPSCs. Mutations in the *HBB* locus result in hemoglobinopathies (e.g., sickle cell disease) which are the most common genetic disorder in the world. The *CCR5* gene encodes a co-receptor in T-cells that is required for infection by the human immunodeficiency virus (HIV). Cell therapies using genome-engineered HSPCs are being developed to treat sickle cell disease, HIV and other hematological disorders.(2,3) Engineered T-cells are used to treat cancer.(4) Similarly, cell therapies derived from HUVECs have been proposed to treat cardiovascular disorders.(27) Recent studies have reported the use of small molecules to inhibit DNA-dependent protein kinase catalytic subunit (DNA-PKcs) and polymerase theta (PolT) to improve the efficiency of gene insertion by blocking NHEJ and MMEJ respectively. We also evaluated the impact of using these compounds on gene insertion using ssDNA with chemical modifications in internal bases (eDNA).(28,29) Finally, we investigated the role of 3’ repair exonuclease 1 (TREX1) in mediating the degradation of unmodified ssDNA.

The results presented below demonstrate that ssDNA templates with chemical modifications in every tenth to every sixth base in addition to the three bases in the end can serve as templates for HR. In addition, eDNA templates show significantly improved gene modification in ABCs, HSPCs, T cells, and HUVECs compared to unmodified ssDNA and ssDNA with CTS. We further show that knocking out *TREX1* results in gene insertion by unmodified ssDNA at levels comparable to eDNA. This indicates that ssDNA degradation is mediated by TREX1 and suggests that eDNA templates resist the activity of this nuclease. Overall, we present a broadly applicable non-viral method to improve genome engineering of multiple clinically relevant cell types.

## Materials and Methods

### Reagents

#### Cells

Human airway basal stem cells (ABCs) were obtained from the Epithelial Cell Core at Nationwide Children’s Hospital (Columbus OH). Other primary cells were obtained commercially. HSPCs were obtained from StemCell Technologies (Catalog: 70060.1) and the Hematopoietic Cell Procurement and Resource Development Center at The Fred Hutchison Cancer Center. HUVECs were obtained from the American Type Culture Collection (ATCC; Catalog: PCS100010). Pan-T cells were obtained from StemCell Technologies (Catalog: 70024).

#### Media

Airway cells: PneumaCult Ex Plus medium (StemCell Technologies #05040) was used for culturing ABCs. Differentiation media was purchased from the University of North Carolina core. HSPC: StemSpan™ SFEM II supplemented with StemSpan™ CD34+ Expansion Supplement (10X) (StemCell Technologies, Catalog # 09720). IPSC: StemFlex™ media (ThermoFisher Scientific, A3349401). HUVEC: EGM™-2 Endothelial Cell Growth Medium-2 BulletKit™ (Lonza, CC-3162). T-cells: X-Vivo 15 media (Lonza; 02-060Q)

#### Small molecules

ROCK inhibitor (Y-27632, MedChemExpress # HY-10583). AZD-7648 (MedChem Express, HY-111783). ART558 (MedChem Express, HY-141520).

#### Antibodies

Anti-Cytokeratin 5 antibody [EP1601Y] (Alexa Fluor® 647) (Abcam, ab193895). Rabbit IgG, monoclonal [EPR25A] -Isotype Control (Alexa Fluor® 647) (Abcam, ab199093). Purified Mouse IgG2b k Isotype Ctrl (Biolegend, 401202). Purified anti-TP63 (W15093A) (Biolegend, 687202). Mouse anti-ACT (Sigma-Aldrich; T6793). Rabbit anti-MUC5B (Sigma-Aldrich; HPA008246). DAPI (Sigma-Aldrich; D9542). AF488 Donkey anti-rabbit (Life Technologies; A21206).

#### Guide RNAs targeting CFTR, HBB, CCR5, and TREX1

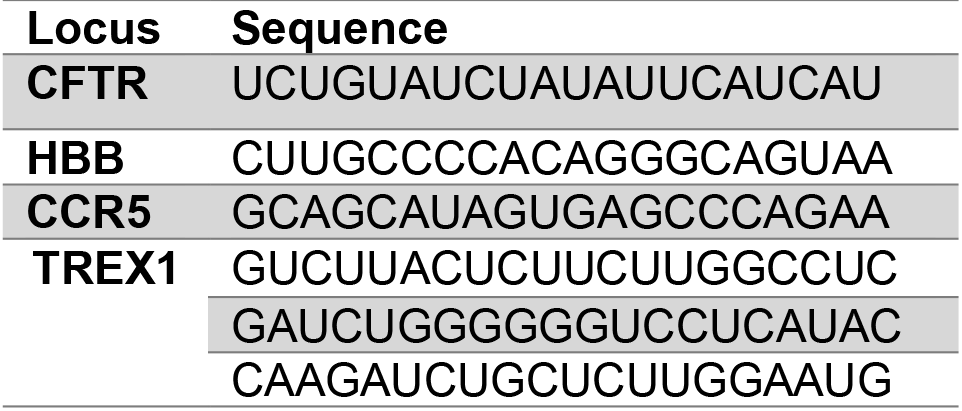

Single guide RNAs were purchased from Synthego Inc. (Redwood city, CA). SgRNAs were chemically modified using 2’O-methyl and phosphorothioate groups in the last three bases.

#### HR Template Sequences

Below are the sequences corresponding to the HR templates for the *CFTR, HBB and CCR5* loci. The lower-case letters correspond to bases mutated relative to the wild type. They were purchased from Integrated DNA Technologies (Coralville, IA).

#### F508del correction sequence(5)

ATGGGAGAACTGGAGCCTTCAGAGGGTAAAATTAAGCACAGTGGAAGAATTTCATTCTGTTCTCAGTTTTCCTGGATTATGCCTGGCACCATTAAAGAAAATATCATCTTcGGcGTgTCtTAcGAcGAgTAcAGATACAGAAGCGTCATCAAAGCATGCCAACTAGAAGAGGTAAGAAACTATGTGAAAACTTTTTGATT

#### G551D sequence – Template 1

AAAAGTGACTCTCTAATTTTCTATTTTTGGTAATAGGACATCTCCAAGTTTGCAGAGAAAGACAATATAGTTCTTGGAGAAGGTGGAATCACAtTatccGGtGacCAACGAGCAAGAATTTCTTTAGCAAGGTGAATAACTAATTATTGGTCTAGCAAGCATTTGCTGTAAATGTCATTCATGTAAAAAAATTACAGAC

#### G551D sequence – Template 2

TAAAAGTGACTCTCTAATTTTCTATTTTTGGTAATAGGACATCTCCAAGTTTGCAGAGAAAGACAATATAGTcttaGGtGAgGGTGGAATCACAtTatccGGtGacCAACGAGCAAGAATTTCTTTAGCAAGGTGAATAACTAATTATTGGTCTAGCAAGCATTTGCTGTAAATGTCATTCATGTAAAAAAATTACAGAC

#### HBB (E6V) correction sequence(3)

CAGGGCTGGGCATAAAAGTCAGGGCAGAGCCATCTATTGCTTACATTTGCTTCTGACACAACTGTGTTCACTAGCAACCTCAAACAGACACCATGGTGCAcCTGACTCCTGAGGAaAAaTCcGCaGTcACTGCCCTGTGGGGCAAGGTGAACGTGGATGAAGTTGGTGGTGAGGCCCTGGGCAGGTTGGTATCAAGGTTA

#### CCR5 sequence

TCATCCTCATCCTGATAAACTGCAAAAGGCTGAAGAGCATGACTGACATCTACCTGCTCAACCTGGCCATCTCTGACCTGTTTTTCCTTCTTACTGTCCCtTTtTGaGCaCACTAgGCTGCCGCCCAGTGGGACTTTGGAAATACAATGTGTCAACTCTTGACAGGGCTCTATTTTATAGGCTTCTTCTCTGGAATCTTC

#### Quantification of Editing

QuickExtract (QE) DNA Extraction Solution (Biosearch Technologies, QE09050). Q5 High Fidelity 2X Master Mix (New England Biolabs, M0492).

### Methods

#### Generation of ssDNA templated with CTS

CTS templates were designed and assembled as described by Shy et al.(19) Similar to other ssDNA templates, the longer ssDNA template was ordered as Ultramers from Integrated DNA Technologies. The template and oligos were resuspended in IDT buffer. The short oligos were mixed with long ssDNA at 4:1 molar ratio. and then heated to 95C and cooled by 5C per 5 mins (1 C per min) in a thermocycler. The annealed product was at a concentration of 8 µM and was used in experiments at a concentration of 800 nM.

#### Cell Culture of ABCs

Human ABCs were cultured in PneumaCult Ex Plus medium (StemCell Technologies #05040) with addition of ROCK inhibitor 10 µM (Y-27632, MedChemExpress # HY-10583) and iMatrix-511 (Nacalai USA #892021) at 5000-10,000 cells/cm^2^ in tissue culture flasks.

#### Cell Culture of HSPCs

G-CSF Mobilized Human Peripheral Blood CD34+ Cells were purchased from StemCell Technologies (Catalog: 70060.1). Cells were thawed by resuspending in 10 mL of RPMI without FBS. The tube was centrifuged at 300 g for 10 minutes and the supernatant removed. Cells were resuspended in StemSpan™ SFEM II supplemented with StemSpan™ CD34+ Expansion Supplement (10X) (StemCell Technologies, Catalog # 09720). Cells were plated on 24-well non-treated plates at a density of 100,000 cells/mL and incubated at 37°C for 2-3 days.

#### Cell Culture of iPSCs

Normal human induced pluripotent stem cells (Gibco, A18945) were cultured in StemFlex™ media (ThermoFisher Scientific, A3349401) at a density of 6.0 × 10^4^ cells/cm^2^ on plates coated with vitronectin (ThermoFisher Scientific, A31804) and incubated at 37°C for 2 days. The ROCK inhibitor, Y-27632, was added to the media 1 h before the experiment at a concentration of 10 µM. Cells were electroporated 2-3 days after plating. After electroporation, cells were plated and cultured for 3 -4 days after which genomic DNA was extracted.

#### Cell Culture of HUVECs

Primary Umbilical Vein Endothelial Cells were purchased from American Type Culture Collection (ATCC, PCS100010). Cells were cultured in EGM™-2 Endothelial Cell Growth Medium-2 BulletKit™ (Lonza, CC-3162) and plated in T75 culture flask at 5000 cells/ cm^2^ with iMatrix-511 (Nacalai USA #892021).

#### Cell Culture of T-cells

Primary T-cells were purchased from StemCell Technologies (#70024). Cells were cultured in X-Vivo 15 media supplemented with 5% human serum and 10 ng/mL of IL-2. T-cells were activated using Dynabeads™ T-cell activator with CD3/CD28 (Thermofisher Inc., #11161D) for 1-2 days using directions provided by the manufacturer (1:1 bead to cell ratio). Failure to remove beads resulted in reduced cell proliferation. Cells were edited after 4-6 days in culture.

#### Gene Editing of ABCs, iPSCs, and HUVECs

Electroporation was performed 4-5 days after plating. Cells were suspended by treatment with TrypLE™ Express Enzyme (1X), phenol red (ThermoFisher Scientific, 12605036). Cells were suspended in Opti-MEM™ I Reduced Serum Medium (ThermoFisher Scientific, 31985088) at 2.5-10 million cells/mL. Cells were electroporated with Lonza 4D-Nucleofector™ in 16-well Nucleocuvette Strips (Lonza, V4XP-3032). Cas9 at 6µg (Integrated DNA Technologies) and sgRNA at 3.2 µg (Synthego) were complexed at room temperature for 10 minutes before adding 100,000 cells (20 µL/well). Cell mixture was transferred to Lonza strip at 20 µL per well and electroporated using program CA-137 with solution setting P3. Afterward, 80 µL of OptiMEM was added to each well. When editing using AAV, AAV was added after electroporation. ABCs and HUVECs were treated with 10^5^ vg/cell. iPSCs were treated with 10^4^ vg/cell. Cells were then transferred to a 12 or 24-well tissue culture plate. Pneumacult Ex Plus (StemCell Technologies #05040), ROCK inhibitor 10 µM (Y-27632, MedChem Express HY-10583), and iMatrix-511 (Nacalai USA #892021) were added to ABCs and EGM™-2 Endothelial Cell Growth Medium-2 BulletKit™ (Lonza, CC-3162) with iMatrix was added to HUVECs and then incubated in 37°C at 5% O_2_. iPSCs were plated with media described above and incubated at 37°C. AZD-7648 (MedChem Express, HY-111783) was added for DNA-PKcs inhibition at a concentration of 0.5µM. ART558 (MedChem Express, HY-141520) was added for PolT inhibition at a concentration of 5 µM.

#### Gene Editing of HSPCs and T-cells

Hematopoietic stem and progenitor cells were counted after two days and centrifuged at 300 g for 10 minutes. T-cells were counted after 4-6 days in culture. Supernatant was removed and cells were resuspended in P3 Primary Cell Nucleofector® Solution with Supplement 1 (Lonza, V4XP-3032) at 5-15 million cell/mL. Cell mixture was transferred to Lonza strip at 20 µL per well and electroporated using program DZ100 with solution setting P3. 80 µL of StemSpan™ SFEM II supplemented with StemSpan™ CD34+ Expansion Supplement (10X) (StemCell Technologies, Catalog #09720) was added to each well. AAV was added after electroporation, if necessary, at 10^4^ vg/cell. HSPCs were transferred to 24-well non-treated plates at a density of 100,000 cells/mL and incubated at 37°C for 2-3 days. T-cells were cultured at densities of 500,000-1,000,000 cells/mL.

#### Gene Correction Measurement

Approximately 3-4 days after electroporation, genomic DNA was extracted using QuickExtract (QE) DNA Extraction Solution (Biosearch Technologies, QE09050). TrypLE Express Enzyme was added to cells, then after 20 minutes they were centrifuged at 300g for 10 minutes. Supernatant was removed, cells were suspended in OptiMEM and counted. 50-100 thousand cells were taken, re-centrifuged and the pellet was suspended in 50 µL of QE. The QE suspension was heated at 65°C for 6 minutes then heated at 98°C for 10 minutes. Amplification was done with the following primers:

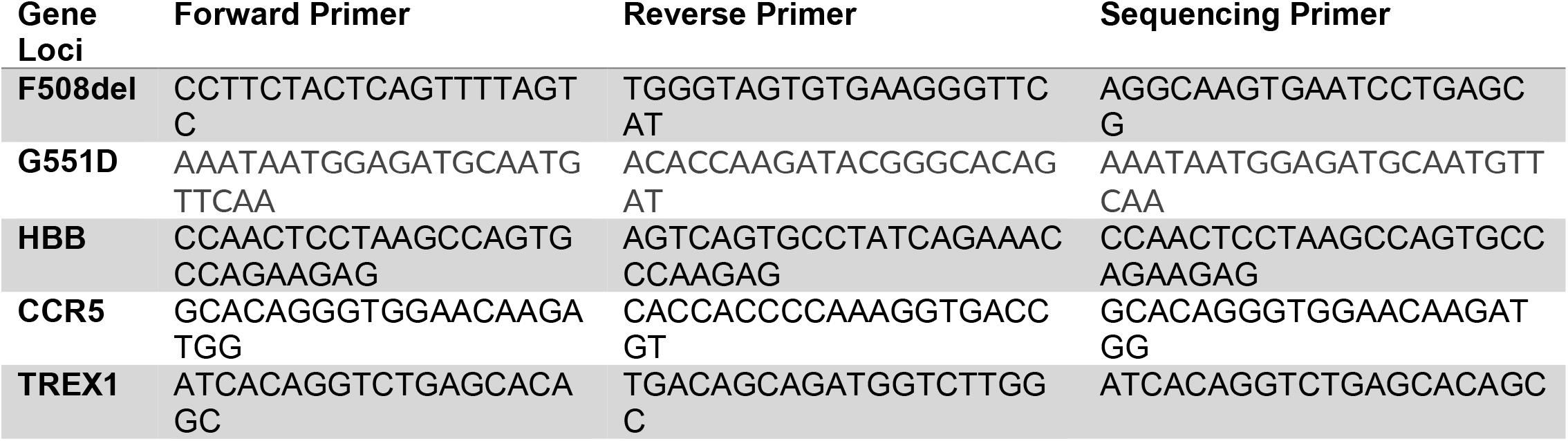

PCR was performed using Q5 High Fidelity 2X Master Mix (New England Biolabs, M0492) with the following temperature and times.

| Gene Site | Temp/<br>minutes | Temp/<br>minutes | Temp/<br>minutes | Temp/<br>minutes | Temp/<br>minutes | Temp/<br>minutes | Temp/<br>minutes |
| --- | --- | --- | --- | --- | --- | --- | --- |
|  | 1 | 2 | 3 | 4 | 5 | 6 | 7 |
|  |  | Denature | Anneal | Extend |  |  |  |
| **F508del** | 98°C/3:00 | 98°C/<br>0:30 | 58°C/0:20 | 72°C/0:35 | Repeat<br>from step 2<br>35X | 72°C/5:00 | 12°C/∞ |
| **G551D** | 98°C/3:00 | 98°C/<br>0:30 | 62°C/0:20 | 72°C/0:20 | Repeat<br>from step 2<br>35X | 72°C/5:00 | 12°C/∞ |
| **HBB** | 95°C/3:00 | 95°C/<br>0:30 | 62°C/0:20 | 72°C/0:15 | Repeat<br>from step 2<br>34X | 72°C/5:00 | 12°C/∞ |
| **CCR5** | 98°C/3:00 | 95°C/<br>0:30 | 61°C/0:20 | 72°C/0:20 | Repeat<br>from step 2<br>34X | 72°C/5:00 | 12°C/∞ |
| **TREX1** | 95°C/3:00 | 95°C/ 0:20 | 67°C/0:20 | 72°C/0:30 | Repeat<br>from step 2<br>34X | 72°C/5:00 | 12°C/∞ |

The PCR product was sanger sequenced then ICE analysis was used to determine the percentage of corrected alleles.

#### Air-Liquid Interface

Gene corrected ABCs were plated on 6.5 mm Transwell plates with 0.4 µm pore polyester membrane insert (StemCell Technologies, 38024) at 40,000-50,000 cells per well. Pnuemacult Ex Plus and ROCK inhibitor (10 µM) was added to both the apical and basal side of insert and iMatrix was added to the apical insert to support expansion. Once cells became confluent, about 3-4 days, media was removed and ALI medium from the University of North Carolina (UNC media) was added to the basal side of the insert.(30) UNC medium was replaced every other day for four weeks.

#### Immunostaining for epithelial cell characterization

CF-donor cells edited with esDNA and differentiated on ALI transwells for a duration of 4 weeks were stained for cytokeratin 5 (KRT5), acetylated tubulin (ACT), and mucin 5B (MUC5B) to identify basal, ciliated, and secretory cells, respectively. The cells were washed with F-12 (1X) Nutrient Mixture (Ham) (Gibco; 11765054) for 10 minutes on ice. The ALIs were fixed in a solution of 3% sucrose and 4% paraformaldehyde in PBS for a duration of 20 minutes on ice and then washed with PBS. Mouse anti-ACT (Sigma-Aldrich; T6793) was diluted 1:8000 and Rabbit anti-MUC5B (Sigma-Aldrich; HPA008246) was diluted 1:500 in staining buffer. 200µL of the diluted primary antibodies were added to the ALI transwells and incubated overnight at 4°C. The primary antibodies were removed, ALI transwells were washed with PBS, and 200 µL of secondary antibody was added to the ALI transwells. PE Goat anti-mouse IgG (Biolegend; 405307) was diluted 1:200 and AF488 Donkey anti-rabbit (Life Technologies; A21206) was diluted 1:200 in staining buffer. The ALI transwells were incubated in secondary antibody for a duration of 1 hour. The secondary antibodies were removed, washed with PBS, and Recombinant Alexa Fluor 647 anti-Cytokeratin 5 antibody diluted 1:200 (Abcam, ab193895) and DAPI (Sigma-Aldrich; D9542) diluted 1:1000 in staining buffer were added to the ALI transwells and incubated for 1 hour. The ALI transwells were washed with PBS, placed on slides using Fluoromount-G Mounting Medium (Invitrogen; 00-4958-02), and cover slipped.

#### Microscopy

Transwell slides were imaged on a motorized Eclipse Ti2-E inverted microscope (Nikon Instruments) with a SOLA LED engine (Lumencor) and an ORCA Fusion CMOS camera (Hamamatsu). Semrock filter sets were used for individual DAPI, GFP, TRITC, and Cy5 channels. Multichannel images were captured as Z-stacks using NIS-Elements AR software (Nikon Instruments, v. 5.30) with 40x Plan Apochromat Lambda objectives (Nikon

Instruments) for a final resolution of 0.32 µm/pixel and 0.16 µm/pixel, respectively. The Z-stacks were compressed into 2D images using the Extended Depth of Focus (EDF) module in NIS-Elements.

#### Ussing Chamber Functional Assay

Four weeks after cells were plated in the transwell plates, ion channel function of CFTR was assessed with the Ussing chamber system (VCC MC6 from Physiologic Instruments Inc. San Diego, CA) with the U2500 Self-contained Ussing chambers (Warner Instruments, Hamden CT). To measure Cl^−^ secretion, the fully-differentiated epithelial sheets were placed with mucosal solution on apical side [120mM NaGluconate, 25mM NaHCO_3_, 3.3mM KH_2_PO_4_, 0.8mM K_2_HPO_4_, 4mM Ca(Gluconate)_2_, 1.2mM Mg(Gluconate)_2_, 10mM mannitol] and serosal solution on basal side [120mM NaCl, 25mM NaHCO_3_, 3.3mM KH_2_PO_4_, 0.8mM K_2_HPO_4_, 1.2mM CaCl_2_, 1.2mM MgCl_2_, 10mM Glucose]. Following this, ion channel activators and inhibitors were added as follows: 10µM amiloride apically to inhibit Epithelial Sodium Channel (ENaC), 10µM forskolin bilaterally (Abcam #ab120058) to activate CFTR channels via activation of adenylyl cylase concentrations, 10µM CFTR potentiator drug VX-770 (SelleckChem, #S1144) apically, 10µM CFTR_inh-172_ (Sigma Millipore, #C2992) apically to inhibit CFTR channel, and 100µM uridine-5’-triphosphate trisodium (UTP) salt (Thermo Scientific Chemicals #AAJ63427MC) apically.

#### Staining and Fluorescence Activated Cell Sorting (FACS) Analysis

Cells were stained 3-4 days after editing with anti-Cytokeratin 5 antibody [EP1601Y] (Alexa Fluor® 647) (Abcam, ab193895), Rabbit IgG, monoclonal [EPR25A] -Isotype Control (Alexa Fluor® 647) (Abcam, ab199093), Purified Mouse IgG2b k Isotype Ctrl (Biolegend, 401202), and purified anti-TP63 (W15093A) (Biolegend, 687202). Flow cytometry was done on the LSR/Fortessa cytometer and data was analyzed using FlowJo software (FlowJo, LLC).

## Statistical Analyses

For experiments comparing editing efficiencies using more than two types of ssDNA templates and/or AAV, significance was assessed using One Way Analysis of Variance (ANOVA) followed by multiple comparisons using Tukey’s test. Paired T-test (two-tailed) was performed for results in Figure 3A-E.

## RESULTS

### Extensive chemical modifications improve gene insertion in airway cells

To test our hypothesis that extensive chemical modification of ssDNA may improve gene editing, we used ssDNA templates with phosphorothioate (PS) modifications in the last three bases (end-modified), at every sixth and every third base in addition to the three bases at the ends (Figure 1A-B). We targeted exon 11 in the *CFTR* locus and attempted to incorporate silent mutations in the region associated with the F508del mutation that is commonly seen in people with CF (pwCF) (Figure S1A). We edited non-CF ABCs with ssDNA templates that were either unmodified or chemically modified. Gene editing using ssDNA was inefficient (<10% alleles with HR) when we used ssDNA templates at a concentration of 800 nM in the absence of the DNA-PKcs inhibitor AZD-7648 (Figure S1B) and there was no improvement in gene insertion at higher concentrations of ssDNA (data not shown). In the presence of AZD-7648, ssDNA templates with PS modification in every sixth base had higher HR (41 ± 15%) than both unmodified (15 ± 5%) and end-modified (24 ± 6%) ssDNA templates (Figure 1C). Optimal gene insertion was achieved using ssDNA templates modified at every sixth base at a concentration of 800 nM (Figure S1C). Interestingly, ssDNA templates with PS modification in every third base showed a significant reduction in HR, suggesting an intermediary level of modifications is required to achieve optimal HR. The chemically modified ssDNA did not significantly reduce the proliferation of the edited cells when compared to unedited cells (Figure S1D).

**Figure 1.**
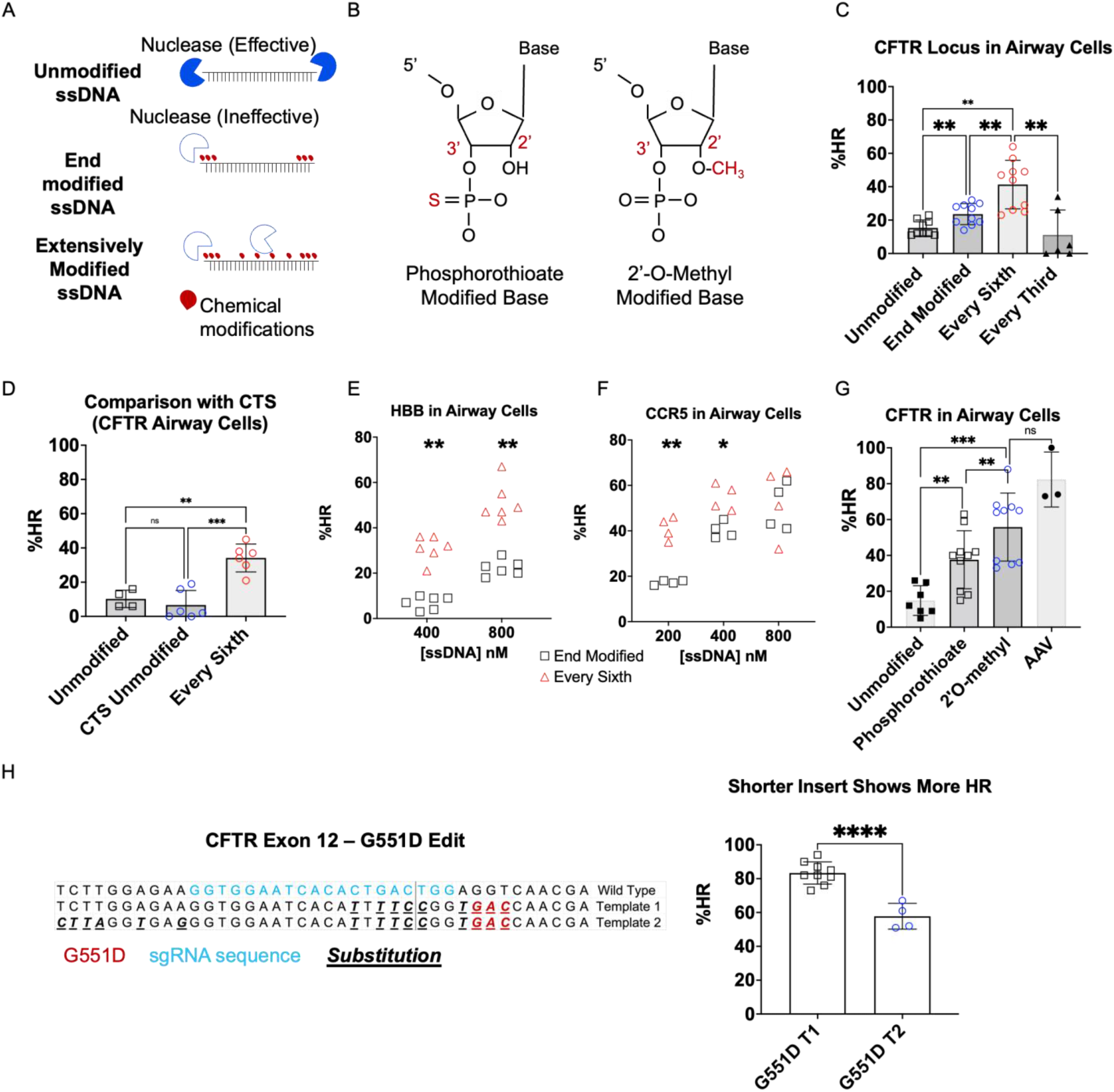
Extensive Chemical Modifications Improve Gene Insertion in Airway Cells. **A**) Graphic showing the chemical modifications of ssDNA and predicted nuclease degradation activity and inactivity. **B**) Graphic showing the chemical structure of modified bases. **C**) In airway cells at the *CFTR* locus, ssDNA templates with every sixth base modified (41 ± 15%) show significantly higher HR than end-modified ssDNA (24 ± 6%), which are both significantly higher than unmodified ssDNA (15 ± 5%). Modifying every third base results in the least gene insertion (11 ± 15%). Statistical comparison was made using one-way ANOVA. **D**) Chemical modification of every sixth base in ssDNA enables improved gene insertion compared to ssDNA with CTS. **E**) ssDNA modified at every sixth base pair (esDNA) is also more efficient than end-modified ssDNA in the *HBB* locus. The esDNA templates achieved gene insertion in 51 ± 9% alleles at the 800nM concentration as opposed to 22 ± 4% for end-modified ssDNA. At 400 nM, esDNA achieved 31 ± 6% gene insertion compared to 7 ± 3% gene insertion achieved by end-modified ssDNA. Statistical comparison was made using two-way ANOVA. **F**) Using esDNA improves gene insertion compared to end-modified ssDNA in the *CCR5* locus at 200 nM (41 ± 5% vs 17 ± 1%) and 400 nM (55 ± 6% vs 40 ± 4%) but not at 800 nM (53 ± 16% vs 51 ± 10%). Statistical comparison was made using two-way ANOVA. **G**) In airway cells, esDNA with OM modification also shows higher HR than unmodified ssDNA (55 ± 19% vs 15 ± 8%) which is not significantly different from HR obtained using AAV templates (82 ± 15%). (n = 5 biological replicates for esDNA with PS, OM modifications and ssDNA. n = 3 biological replicates for AAV). **H**) Gene insertion of different lengths was attempted in exon 12 of the CFTR locus using esDNA templates to introduce the G551D mutation. Editing was performed with DNA-PKcs inhibition. Template 1 (T1) which replaces 12 bp achieved ∼80-90% gene insertion while template 2 (T2) which replaces 34 bp achieved only ∼60 bp gene insertion. Statistical comparison was made using one-way ANOVA. For all panels using one-way ANOVA, multiple comparisons was performed using Tukey’s test. Sidak’s test was used for two-way ANOVA. ****, ***, ** and * represent p <0.0001, p < 0.001, p< 0.01 and p < 0.05 respectively for all panels.

Cas9 Targeting Sequences (CTS) were recently reported to improve gene insertion using ssDNA by improving nuclear localization of the ssDNA.(19) We designed ssDNA templates with CTS sequences targeting exon 11 of CFTR (Figure S1E). Compared to unmodified ssDNA with CTS (7 ± 9%), ssDNA with PS modifications in every sixth base showed significantly improved gene insertion (34 ± 9%) (Figure 1D). Adding CTS sequences to ssDNA with chemical modifications in every sixth base did not improve gene insertion further (Figure S1F).

To test if these findings can be widely applied in other therapeutically relevant loci, we tested our approach in the *HBB* and *CCR5* loci using ssDNA concentrations ranging from 200-800 nM. We introduced silent mutations in the region corresponding to the sickle cell mutation (E6V) in *HBB* and introduced a premature stop codon in the *CCR5* locus. For *HBB*, ssDNA modified in every sixth base (esDNA) showed improved HR at all concentrations (Figure 1E). For *CCR5*, esDNA improved HR at 200 and 400 nM but not 800 nM (Figure 1F). Overall, esDNA showed significantly higher HR at all loci tested in ABCs with DNA-PKcs inhibition (Figure 1C-F). Gene insertion using esDNA templates is also superior to end-modified ssDNA when both DNA-PKcs and PolT are inhibited (Figure S1G-H).

We then investigated if improved editing can be obtained using other chemical modification s. We used esDNA with 2’O-methyl (OM) modifications on approximately every sixth base (Figure 1B). We avoided modifying thymine since the OM bases are RNA. OM modification in esDNA produced even higher HR (55 ± 19%) than the esDNA with PS modification (38 ± 16%). Gene insertion with OM modified esDNA was not significantly different from gene insertion using AAV templates (82 ± 15%) (Figure 1G). Both esDNA formulations had higher HR than unmodified ssDNA templates (15 ± 8%).

To further understand if higher gene insertion levels can be achieved with shorter inserts, we edited exon 12 in the CFTR locus associated with the variant G551D (another common CF causing mutation) with two different esDNA templates (Figure 1H). Template 1 which replaces 12 bases next to the DSB site enabled gene insertion in 80-90% alleles (Figure 1H). Conversely, template 2 which replaces a 34 bases segment yields ∼60% edited alleles (Figure 1H).

### Gene correction of the F508del mutation in CF ABCs restores CFTR function in differentiated airway epithelial cells

To test the ability to restore CFTR function, we began by gene correcting ABCs homozygous for the F508del mutation. Patient-derived CF ABCs were efficiently corrected using esDNA with PS and OM modifications (Figure 2A). In contrast to the non-CF ABCs, there was no significant difference between esDNA templates with PS or OM (47 ± 14% vs 49 ± 14%). We further validated that esDNA did not result in the introduction of unintended missense mutations in the F508del locus using next-generation sequencing. Figure 2B shows results from one CF sample edited using esDNA with PS modifications. Over 90% of alleles with HR (42% of total alleles) showed perfect insertion of the intended sequence. Editing did not induce a reduction in basal cell markers, cytokeratin 5 (KRT5) and tumor protein P63 (P63) (Figure 2C). On average, all conditions showed > 85% cells that were positive for both KRT5 and P63.

**Figure 2.**
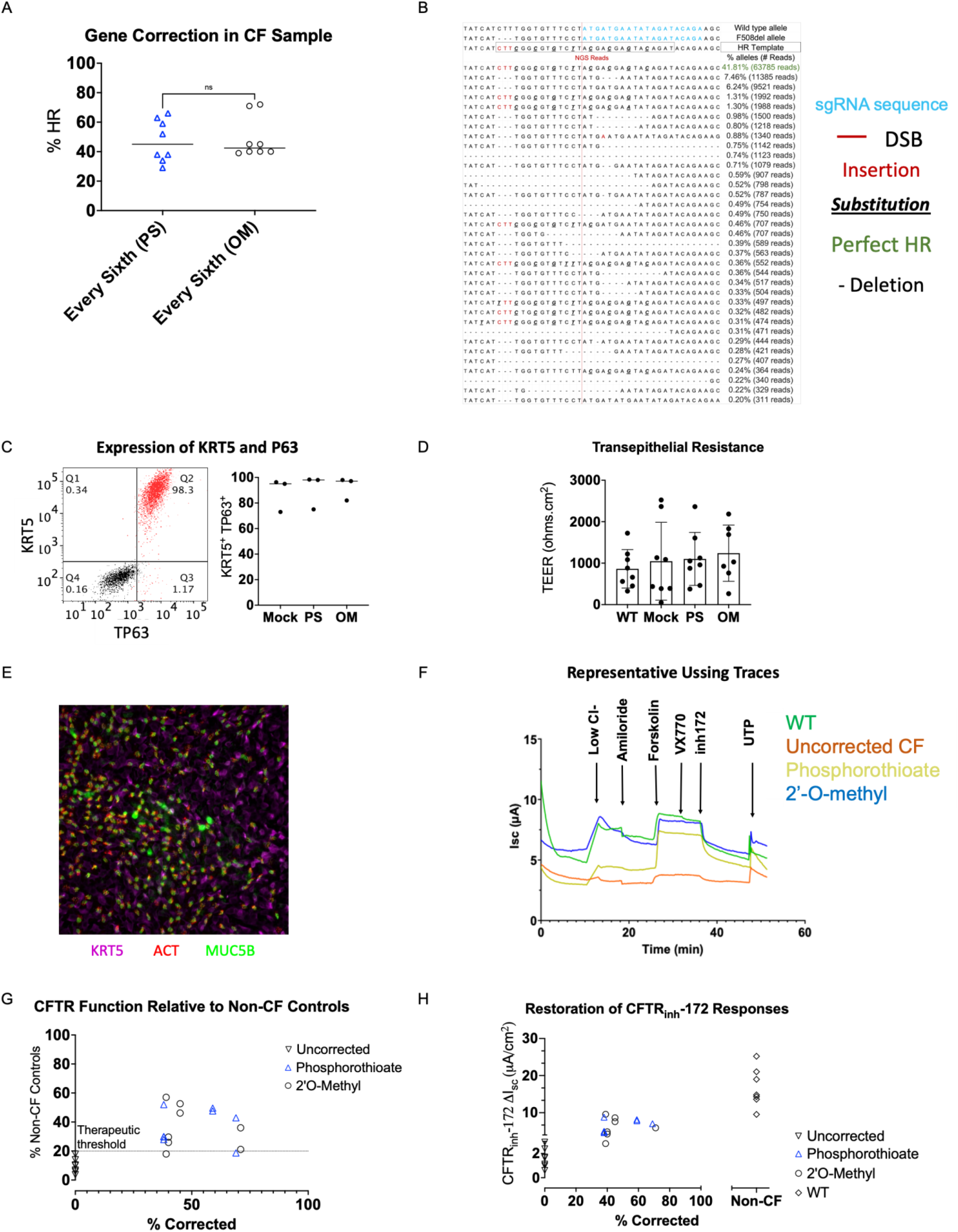
Correction of F508del mutation restores CFTR function in ABCs from pwCF. **A**) Both esDNA OM and esDNA PS corrected ABCs from CF donors (49 ± 14% and 47 ± 14%, respectively). (n=4 biological replicates). **B**) Gene correction of the F508del mutation was confirmed using next-generation sequencing. NGS showed 41.8% alleles with perfect HR and ∼5% alleles with imperfect HR. Unintended substitutions were found in only 0.6% of alleles. The sgRNA sequence is highlighted in blue. **C**) A representative FACS plot shows that > 98% of ABCs were positive for KRT5 and P63 4 days after editing. There was no reduction in KRT5 and P63 expression in the edited ABCs relative to Mock electroporated controls in experiments from three different donors. **D**) Edited samples displayed transepithelial resistances (TEER) similar to unedited controls (WT and Mock), indicating that the editing process does not compromise the ability of the ABCs to produce fully differentiated epithelial sheets. **E)** Edited ABCs produced epithelial sheets containing basal (KRT5), ciliated (ACT) and secretory cells (MUC5B), further indicating the formation of differentiated epithelial sheets. **F**) Representative traces from Ussing chamber analysis showing a non-CF control, an uncorrected CF sample, esDNA PS and esDNA OM corrected CF samples. Uncorrected CF samples show minimal responses to forskolin and CFTR_inh_-172 which activate and inhibit CFTR, respectively. All edited samples showed restored responses to forskolin and CFTR_inh_-172 indicating restored CFTR function. **G**) Percent restoration of CFTR function in corrected CF samples relative to non-CF controls plotted as a function of allelic correction. Edited samples showed 30-60% restoration of CFTR function on average. **H**) Raw CFTR_inh_-172 short circuit currents observed in CF donor ABCs compared to non-CF controls and uncorrected CF controls (n=3 biological replicates). Samples corrected using esDNA show increased response to CFTR_inh_-172 relative to uncorrected controls. This change in current was statistically significant when compared using one-way analysis of variance (ANOVA) followed by multiple comparisons test (p < 0.05).

To quantify the function of CFTR following gene correction, corrected CF, uncorrected CF, and non-CF samples, were grown at air liquid interface (ALI) to induce differentiation into polarized airway epithelial sheets and then currents were measured using Ussing chamber assay. Epithelial barrier integrity was evaluated by measuring the transepithelial resistance (TEER). Edited ABCs produced epithelial sheets with TEER comparable to controls thus indicating that the edited ABCs can form differentiated epithelial sheets (Figure 2D). ALI cultures contained basal cells expressing KRT5, ciliated cells expressing acetylated tubulin (ACT) and secretory cells producing mucin 5B (MUC5B) (Figure 2E). The corrected CF samples showed restored CFTR function indicated by increased forskolin-stimulated currents and increased CFTR_inh-_172-sensitive short circuit current (Figure 2F-H). On average, uncorrected CF samples showed 1.6 ± 0.9 µA/cm^2^ change in short circuit currents in response to CFTR_inh_-172 compared to 16 ± 6 µA/cm^2^ for non-CF controls. CF samples corrected using PS and OM esDNA showed 6 ± 2 µA/cm^2^ change in short circuit currents in response to CFTR_inh_-172. This change in current was statistically significant when compared using one-way analysis of variance (ANOVA) followed by multiple comparisons test (p < 0.05). When we compared CFTR function in gene corrected cells to non-CF controls, epithelial sheets derived from ABCs edited using esDNA with PS and OM showed CFTR function above the therapeutic threshold (20%) for clinical benefit (37 ± 12% for esDNA with PS and 36 ± 15% for esDNA with OM relative to non-CF controls) (Figure 2G).

### Extensive chemical modification in ssDNA templates improves gene insertion in HSPCs, T-cells, and HUVECs but not iPSCs

To further test if esDNA would be effective in improving gene insertion in other therapeutically relevant primary cells, we tested them in HSPCs, HUVECs, T-cells, and iPSCs. In HSPCs, we tested editing in the *HBB, CFTR*, and *CCR5* loci and found that esDNA improved HR in all three loci. In HSPCs, esDNA (47 ± 22%) improved HR over end-modified ssDNA (18 ± 13%) in the *HBB* locus (Figure 3A). Similar improvement in gene insertion was seen in *CFTR* (20 ± 6 % vs 5 ± 5 % for esDNA vs end-modified) and *CCR5* (38 ± 26% vs 15 ± 23% for esDNA vs end-modified) loci in HSPCs (Figures 3B-C). In HSPCs, gene insertion by esDNA templates was also more efficient compared to end-modified ssDNA in the absence of DNA-PKcs inhibition (Figure S2A). We tested esDNA in HUVECs and T-cells by adding silent mutations in the *CFTR* locus. In HUVECs, the use of esDNA resulted in 12 ± 5% alleles with HR compared to 4 ± 3% alleles with HR obtained with end-modified ssDNA (Figure 3D). In T-cells, esDNA templates resulted in 30 ± 10% compared to 13 ± 6% obtained with end-modified ssDNA (Figure 3E).

**Figure 3.**
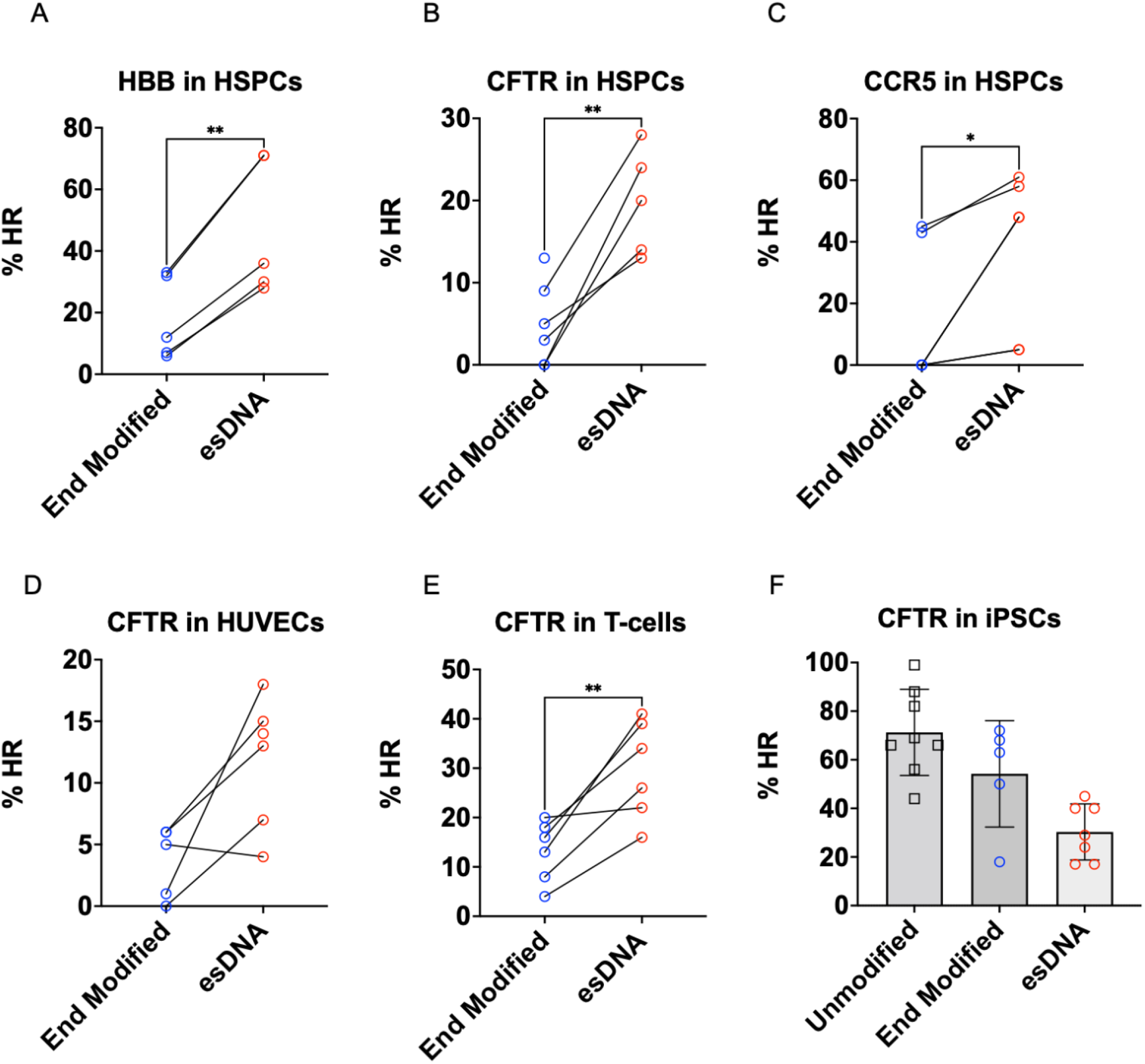
esDNA templates improve gene insertion in HSPCs, T-cells and HUVECs but not iPSCs. HR in HSPCs significantly increased with esDNA compared to end-modified ssDNA in the **A**) *HBB* locus (47 ± 22% vs 18 ± 13%), **B**) *CFTR* locus (20 ± 6% vs. 5 ± 5%), and **C**) *CCR5* locus (38 ± 26% vs. 15 ± 3%). **D**) In HUVECs, esDNA increased HR in the CFTR locus from 4 ± 3% obtained with end-modified ssDNA to 12 ± 5% (p = 0.0531). **E**) In T-cells, esDNA increased HR in the CFTR locus from 13 ± 6% obtained with end-modified ssDNA to 30 ± 10%. **F**) Gene insertion with esDNA (30 ± 12%) was significantly lower in iPSCs in the *CFTR* locus than unmodified ssDNA (71 ± 17%). 3 biological replicates were tested for each cell type. Groups in A-E were compared using a paired T-test. ** and * represent p <0.01, p < and p < 0.05 respectively for all panels.

We also tried adding silent mutations in the *CFTR* locus in iPSCs using esDNA. Surprisingly, we found that in iPSCs, unmodified ssDNA was more effective in enabling HR-mediated gene insertion over end-modified ssDNA (71 ± 18% vs 54 ± 22%). Conversely, esDNA showed reduced HR (30 ± 12%) compared to all other conditions (Figure 3F). Similar results were observed when iPSCs were edited in the absence of DNA-PKcs inhibition (Figure S2B).

### Knocking out TREX1 exonuclease improves HR using unmodified ssDNA templates

We were interested in characterizing the mechanism by which extensive chemical modifications protect ssDNA. We reasoned that the inactivation of one or more of these nucleases may enable improved gene insertion by ssDNA. A recent pre-print reported that the nuclease TREX1 may be involved in degrading ssDNA in cell lines.(31) To investigate the effect of TREX1 on ssDNA, we genome edited ABCs to knock out *TREX1* (TREX1 KO) by using Cas9 to create insertions and deletions in the gene (Figure 4A). HR using unmodified ssDNA was dramatically improved in *TREX1* KO ABCs compared to wild type (WT) (4 ± 4% for WT and 44 ± 14% for TREX1-KO). However, HR was not improved in *TREX1* KO ABCs when esDNA template was used (46 ± 4% for WT and 41 ± 11% for TREX1 KO) (Figure 4B). HUVECs and T-cells with *TREX1* KO also showed improved gene insertion using unmodified ssDNA (Figure S3A-B). These studies suggest that extensive chemical modifications protect esDNA from TREX1.

**Figure 4.**
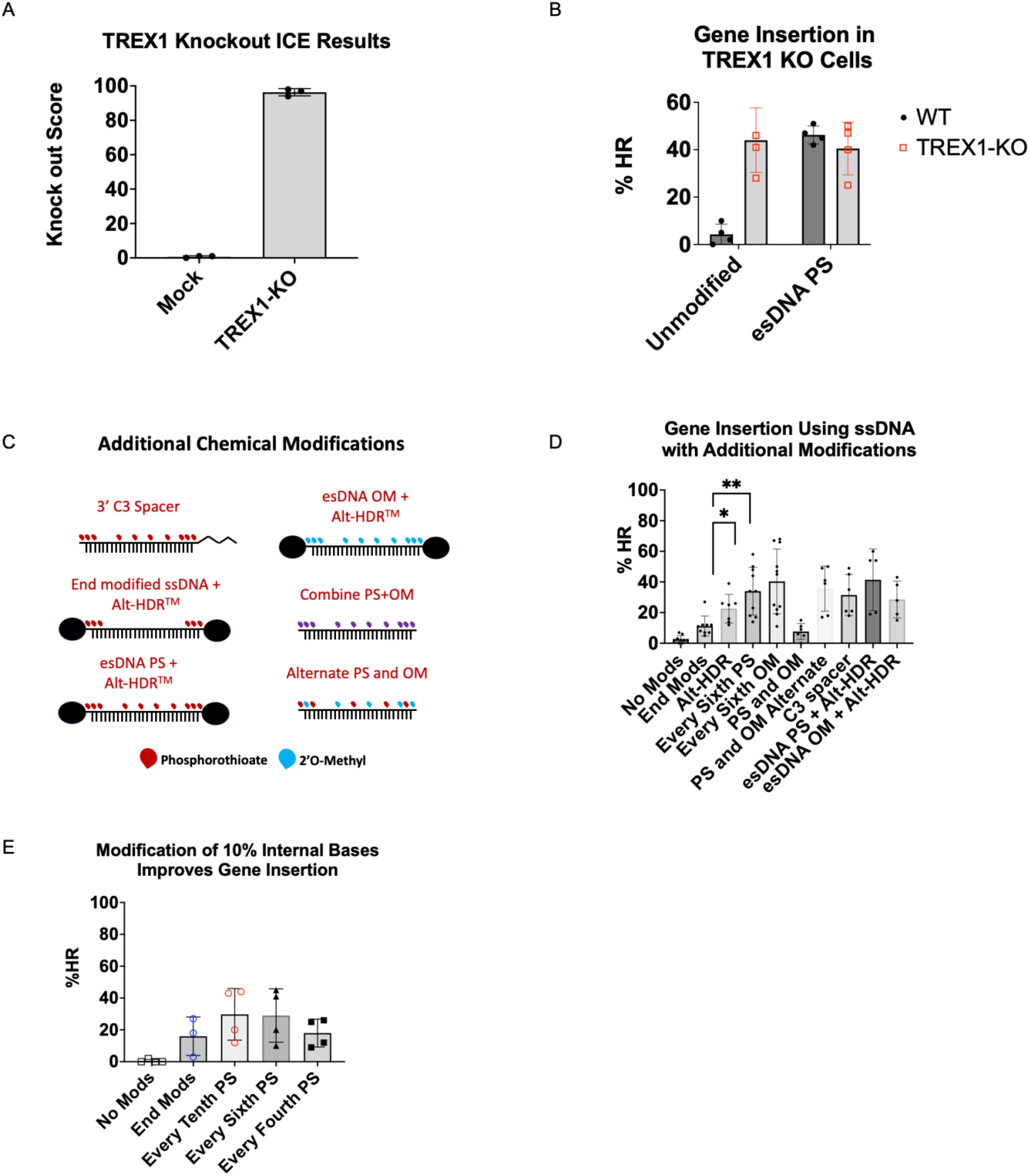
Knocking out *TREX1* improved Gene Insertion without chemically modifying ssDNA. **A**) Percentage of alleles with *TREX1* knocked out in non-CF ABCs samples (n=3 biological replicates). **B**) HR using unmodified ssDNA templates was improved when *TREX1* was knocked out in non-CF ABCs. Knocking out *TREX1* did not improve HR by esDNA templates (n = 4 biological replicates). **C**) Graphic showing additional chemical modification combinations including commercially available proprietary end modifications. **D**) Gene insertion with ssDNA using combinations of chemical modifications in ABCs. Both esDNA OM (40 ± 21%) and esDNA PS (34 ± 16%) showed improved gene insertion relative to unmodified (3 ± 3%), end-modified (11 ± 6%) and end-blocked ssDNA (Alt-HDR) (23 ± 9%). All other combinations did not improve on esDNA OM (40 ± 21%) and esDNA PS (34 ± 16%) (n=6 biological replicates). **E**) Modification of every 10^th^ base with PS groups also improved gene insertion compared to unmodified or end-modified ssDNA. Gene insertion was reduced when every fourth base was modified. ** and * represent p <0.01, p < and p < 0.05 respectively in all panels.

Since modification of the 3’ end has been reported to protect ssDNA from TREX1,(31) we tested if protection of 3’ end by modifying the OH group in esDNA would improve gene insertion further (Figure 4C). The addition of a 3’ C3 spacer to esDNA did not improve gene insertion (Figure 4D). We also investigated if proprietary end protecting modifications available from Integrated DNA Technologies (IDT) (Alt-HDR modifications) may enhance HR further. Alt-HDR modifications added to end-modified ssDNA improved gene insertion relative to end modification with PS alone (Figure 4D). The use of esDNA still showed higher gene insertion relative to Alt-HDR modifications alone. However, there was no further improvement in gene insertion when Alt-HDR modifications were added to esDNA. We reasoned that increased protection of ssDNA by combining modifications may slow degradation by ssDNA further. Therefore, we investigated if using templates with combinations of PS and OM modifications improved gene insertion further (Figure 4C). However, this was not the case (Figure 4D). Since modification of every third base reduced gene insertion, we wanted to assess if modifying fewer internal bases than every sixth would further improve gene insertion. Modifying every tenth base in ssDNA resulted in 30 ± 16% alleles with HR which was similar to esDNA (29 ± 17%) (Figure 4E). Modifying every fourth base reduced alleles with HR to 18 ± 9%.

We wanted to further validate that the activity of TREX1 was specific to ssDNA. Therefore, we tested the impact of TREX1 on AAV-based HR templates targeting the *CFTR* locus.(5) These AAV templates have significantly longer homology arms (>800 bp) compared to the ssDNA and esDNA templates (∼90 bp) (Figure S3C). AAV-based HR templates can be used to reliably edit the *CFTR* locus in ABCs, HSPCs, HUVECs and iPSCs (Figure S3D). When using DNA-PKcs inhibition, we obtained ∼90% modified alleles in airway cells and 60-80% modification in HSPCs using AAV-based templates (Figure S3D). ABCs with *TREX1* KO did not show any change in gene insertion of the *CFTR* locus when edited using Cas9 and AAV-based templates in the presence of DNA-PKcs inhibition (Figure S3E). Thus, the activity of TREX1 is specific to ssDNA.

## Discussion

Previous studies investigating the use of ssDNA to perform HR-mediated gene insertion have explored the protection of the last 3 bases of the ssDNA templates but not internal bases.(8,17-19) Although chemical modification of internal bases has been used to protect sgRNA and mRNA used in gene editing, the approach may not have been an obvious choice for HR templates due to differences in the biological processes involved.(20,21,24) In the case of sgRNA, the modified bases do not interact with host proteins. Thus, modification of sgRNA has a lower chance of compromising nuclease activity as long as the modifications don’t hinder base pairing or interactions with the nuclease. In the case of mRNA, many of the most successful modifications are already present in nature and thus compatible with the natural translational apparatus in cells. By way of contrast, chemical modifications in HR templates must be compatible with a variety of proteins involved in the HR pathway. Our results show that the HR machinery can indeed process chemically modified bases. Our results also show that this compatibility with the HR machinery can be lost if too many bases are modified. We only tested two types of chemical modifications. It is possible that other types of chemical modifications may yield similar results. The combination of at least the OM and PS modifications did not improve editing further. It is unclear if combining other base modifications with PS or OM bases will improve gene insertion further.

Surprisingly, the modified bases do not appear to compromise the fidelity of HR. The next generation sequencing results show that the chemical modifications are processed similarly to normal bases by the HR machinery since significant unintended missense mutations were not present in the edited cells. Furthermore, CF airway cells corrected using esDNA templates show restored CFTR function (Figure 2F-H). Further studies may be needed to assess the ability of esDNA templates to correct disease-associated phenotypes in other cell types and to investigate if other modifications affect the fidelity of HR.

AAV-based templates, which have significantly longer homology arms (>800 bp), can consistently achieve HR in up to 90% of alleles in the presence of DNA-PKcs inhibition (Figure 1G, Figure 3SD). Our results show that the use of the right modification (Figure 1G) and design (Figure 1H) allows the use of eDNA to modify a similar proportion of alleles as AAV templates (80-90% of alleles). It has previously been shown that gene insertion in the *CFTR* locus in airway cells can be increased by ∼2-fold by increasing homology arm length to >150 bp.(32) We posit that long ssDNA with chemical modifications in 10-17% of internal bases may show even better gene insertion. However, such long chemically modified ssDNA templates are not available commercially and are challenging to produce.

The use of eDNA templates to achieve consistent gene insertion relies heavily on the availability of effective DNA-PKcs inhibitors that improve HR by blocking NHEJ.(28) We observed efficient gene insertion in ABCs using ssDNA templates only in the presence of the DNA-PKcs inhibitor, AZD 7648. We observed measurable HR in iPSCs and HSPCs without DNA-PKcs inhibition and the use of esDNA still improved gene insertion relative to ssDNA in HSPCs (Figure S2A). Thus, although extensive chemical modification of ssDNA improves gene editing in multiple cell types, the need for DNA-PKcs inhibition to obtain detectable gene insertion using ssDNA or esDNA appears to be cell-type dependent. It is possible that the differences in the efficiency of gene insertion correlate with the degradation of ssDNA in these cells. This assumption is driven by the fact that AAV-based templates enable similar gene modification in all the cell types tested (Figure S3D).

In attempting to identify nucleases that may be involved in ssDNA degradation, we investigated TREX1 primarily because *TREX1* KO was recently reported to improve gene insertion using ssDNA templates in multiple immortalized cell lines.(31) Our study validates this observation in three primary cell types and shows that knocking out *TREX1* can improve gene insertion using unmodified ssDNA templates. Moreover, since knocking out *TREX1* did not improve gene insertion using AAV (Figure S3E), our results further show that the role of TREX1 in genome editing is specific to ssDNA. It has already been proposed that protecting the ends increases HR by preventing TREX1 from degrading ssDNA.(31) However, our data shows that modifying internal bases along with the ends somehow further improves gene insertion. Notably, our study also shows that the knockout of TREX1 can protect the ssDNA as efficiently as chemical modifications.

The esDNA templates described here may be extremely useful in characterizing the biological impact of small INDELS and missense mutations. Previous studies have attempted to model the biology of CFTR variants by introducing them into cell lines in vitro.(14) Such models are valuable since they are isogenic and account for the impact of modifier genes that results in widely varying disease phenotypes.(33) Similarly, Cas9 and ssDNA templates have been used to model the role of mutations in TP53 in myeloid malignancies in cell lines such as K562.(34) Attempts have also been made to model oncogenic mutations in other genes in HSPCs and epithelial stem cells directly.(15,35) The use of eDNA templates with DNA-PKcs inhibition may thus enable efficient generations of primary cells with such variants and aid such studies significantly.

Apart from modeling the impact of mutations, cell therapies have been proposed to treat genetic disorders caused by missense mutations and small INDELs such as sickle cell disease and CF.(3,5) The ∼40% allelic correction of the F508del mutation reported here enables the partial restoration of CFTR function (∼35% of non-CF) in airway cells obtained from people with CF. This is therapeutically relevant because restoration of ∼20% CFTR function relative to non-CF controls is thought to be sufficient to provide therapeutic benefit.(36,37) Similarly, correction of 20-60% alleles in HSPCs from donors with sickle cell disease has been shown to restore mature hemoglobin production and prevent sickling.(38) One advantage of short eDNA templates is that they can be readily produced in a few days as opposed to AAV template construction and packaging, which can take a few weeks to months. In addition, they may not elicit the innate immune responses which have been reported to compromise the regenerative potential of HSPCs edited using Cas9 and AAV.(8,12)

Apart from biological reasons, non-viral templates that can be mass produced readily are also of interest for improving access to genetic therapies globally.(39) Personalized ex-vivo genetic therapies require complex manufacturing of cells and AAV. In addition, they also require complex medical procedures that are not feasible in low-resource environments. In vivo gene editing using off-the-shelf reagents may enable widespread gene correction to treat genetic diseases globally.(39) Notably, in vivo gene editing using lipid nanoparticles packaged with Cas9 mRNA, sgRNA and ssDNA has been proposed to correct CFTR mutations in vivo.(40) Future studies may test the use of eDNA packaged in LNP for correcting pathogenic mutations using in vivo gene editing. Non-viral in vivo editing approaches are less likely to be limited by immune responses which limit the efficacy of viral vectors such as AAV. LNPs targeting organs such as muscles which were traditionally only targetable using AAV have also been recently reported.(41) Thus, the findings presented here can have broad therapeutic implications.

Other non-viral templates have been reported for gene insertion including ssDNA with CTS and double-stranded DNA with interstrand cross-linking.(19,42) We did not see improved gene insertion using CTS in airway cells. It is possible that the observation may be due to the cell type or locus targeted. Compared to the CTS approach, eDNA does not require the design of special sequences and can thus be applied readily without additional effort or room for errors. Interstrand crosslinked dsDNA has not been reported for use in stem cells such as HSPCs and may be limited by innate immune responses that have been previously shown to limit gene insertion using dsDNA templates in primary cells.(43) Although further studies are needed to validate the impact of eDNA templates on the regenerative potential of ABCs and HSPCs, the results presented here represent a starting point for further improving the efficiency of ssDNA templates using internal chemical modifications.

## Conclusion

Our study shows that the homologous recombination machinery in primary cells can tolerate modified bases in the template. The results show that chemical modification of internal bases in ssDNA templates at every tenth to every sixth base significantly improves gene insertion using CRISPR-Cas9 in multiple clinically relevant human donor-derived primary cell types (HSPCs, T-cells, ABCs and HUVECs). Upon optimization, gene insertion can be achieved in 80-90% of alleles which is comparable to AAV mediated gene insertion. In addition, our study shows that the chemical modifications work by protecting the ssDNA from degradation by TREX1. Overall, we present a broadly applicable method to improve the insertion of short sequences in primary cells using CRISPR-Cas9 and chemically modified ssDNA templates. These tools can enable improved production of cell therapies and enable studies characterizing the biological impact of different genetic variants in primary cells.

## Data Availability

Raw data is available at : https://zenodo.org/doi/10.5281/zenodo.10402324

## Acknowledgments

We thank Dr. Matthew Porteus from Stanford University for sharing HR template plasmids targeting the F508del mutation. Source lung tissues were provided to ECC by the Comprehensive Transplant Center Human Tissue Biorepository of The OSU Wexner Medical Center. Some HSPCs samples were obtained from the Cooperative Centers of Excellence in Hematology NIDDK at the Fred Hutchison Cancer Center supported by the National Institute of Diabetes, Digestive and Kidney Diseases (Grant #: DK106829).

