## Supplementary Data for "Single-Stranded DNA with Internal Base Modifications Mediates Highly Efficient Gene Insertion in Primary Cells"

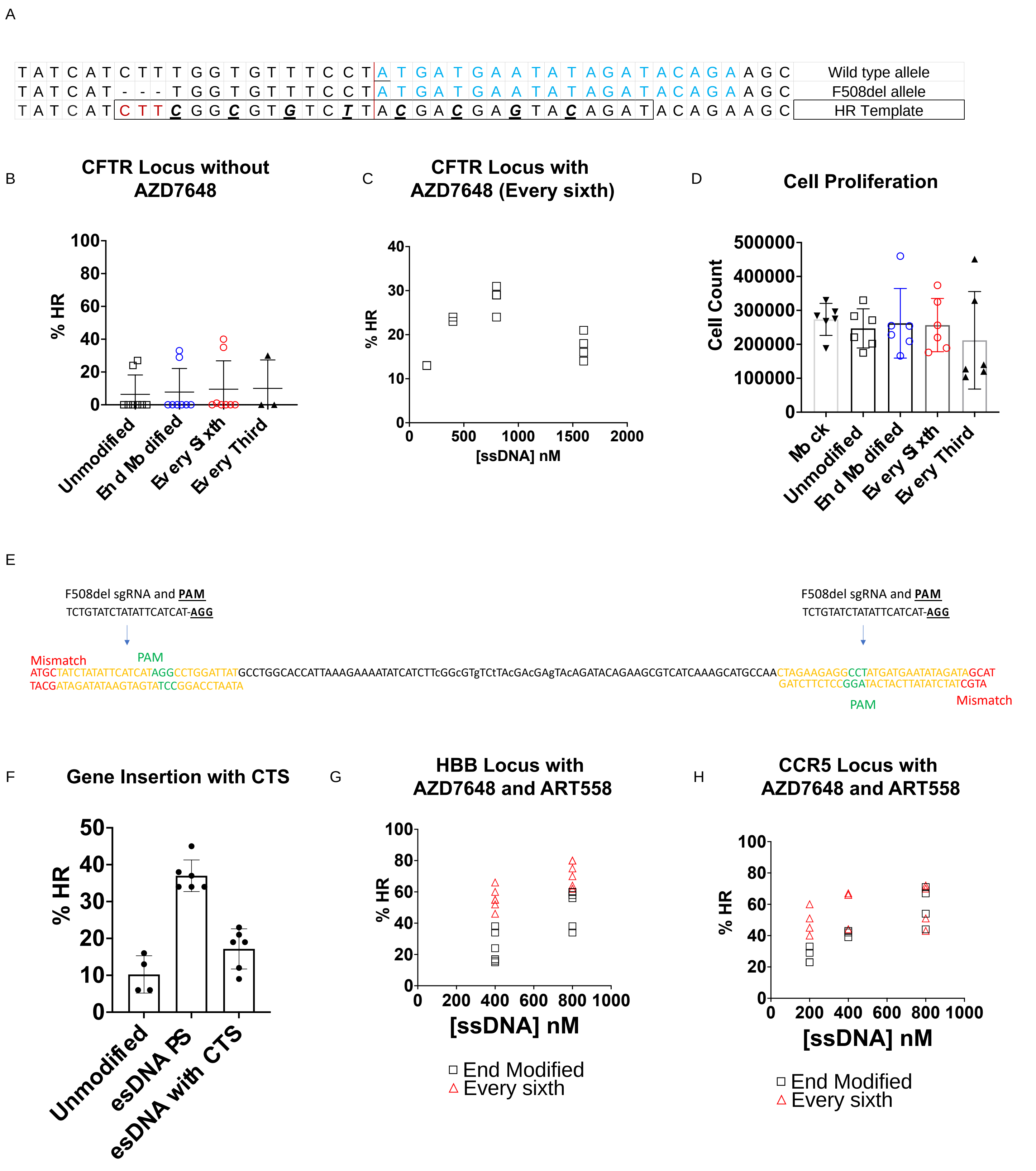


**Figure S1**: Role of NHEJ and MMEJ inhibition on gene insertion using ssDNA and esDNA.

**A)** Graphic comparing the F508del allele with the wild-type *CFTR* allele and HR template. F508del involves the deletion of the sequence CTT in exon 11 of CFTR. The HR template includes the missing bases and additional silent mutations to prevent recutting by Cas9. The sgRNA sequence is highlighted in blue. CTT is highlighted in red in the HR template, and the silent mutations are bolded, underlined and italicized. **B**) Gene insertion using ssDNA was inefficient in airway cells in the absence of DNA-PKcs inhibitor (n = 4 biological replicates). **C**) In the presence of a DNA-PKcs inhibitor, gene insertion in the CFTR locus was optimal at an esDNA concentration of 800 nM. **D**) Gene insertion using ssDNA templates does not reduce cell proliferation significantly. **E**) Cas9 targeting sequences which have been reported to help localize ssDNA templates to the nucleus were added to esDNA templates. Graphic describing the design of esDNA templates with CTS sequences **F**) There was no improvement in gene insertion by esDNA templates in the presence of CTS sequences. The use of DNA-PKcs inhibitor and polymerase theta inhibitor (ART558) enables the use of lower concentrations of esDNA to achieve improved gene insertion in the **G**) *HBB* locus and **H**) *CCR5* locus (n = 2-3 biological replicates). However, esDNA was still superior to end-modified ssDNA.

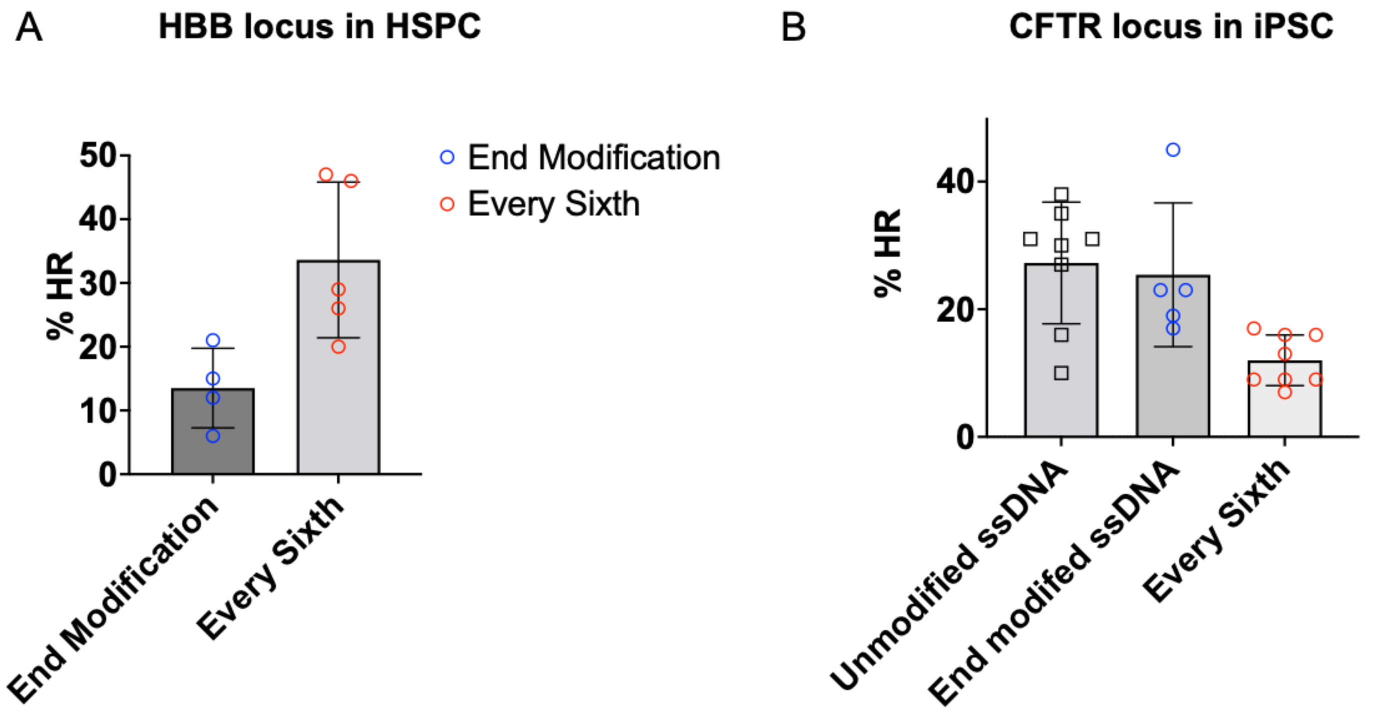


**Figure S2**: Gene insertion using ssDNA and esDNA in HSPCs and iPSCs in the absence of DNA-PKcs inhibition.

**A**) Gene insertion was superior using esDNA templates in HSPCs even in the absence of DNA-PKcs inhibition (n = 3 biological replicates). **B**) Unmodified ssDNA was sufficient for gene insertion in iPSCs even in the absence of DNA-PKcs inhibition. (n = 3 biological replicates).

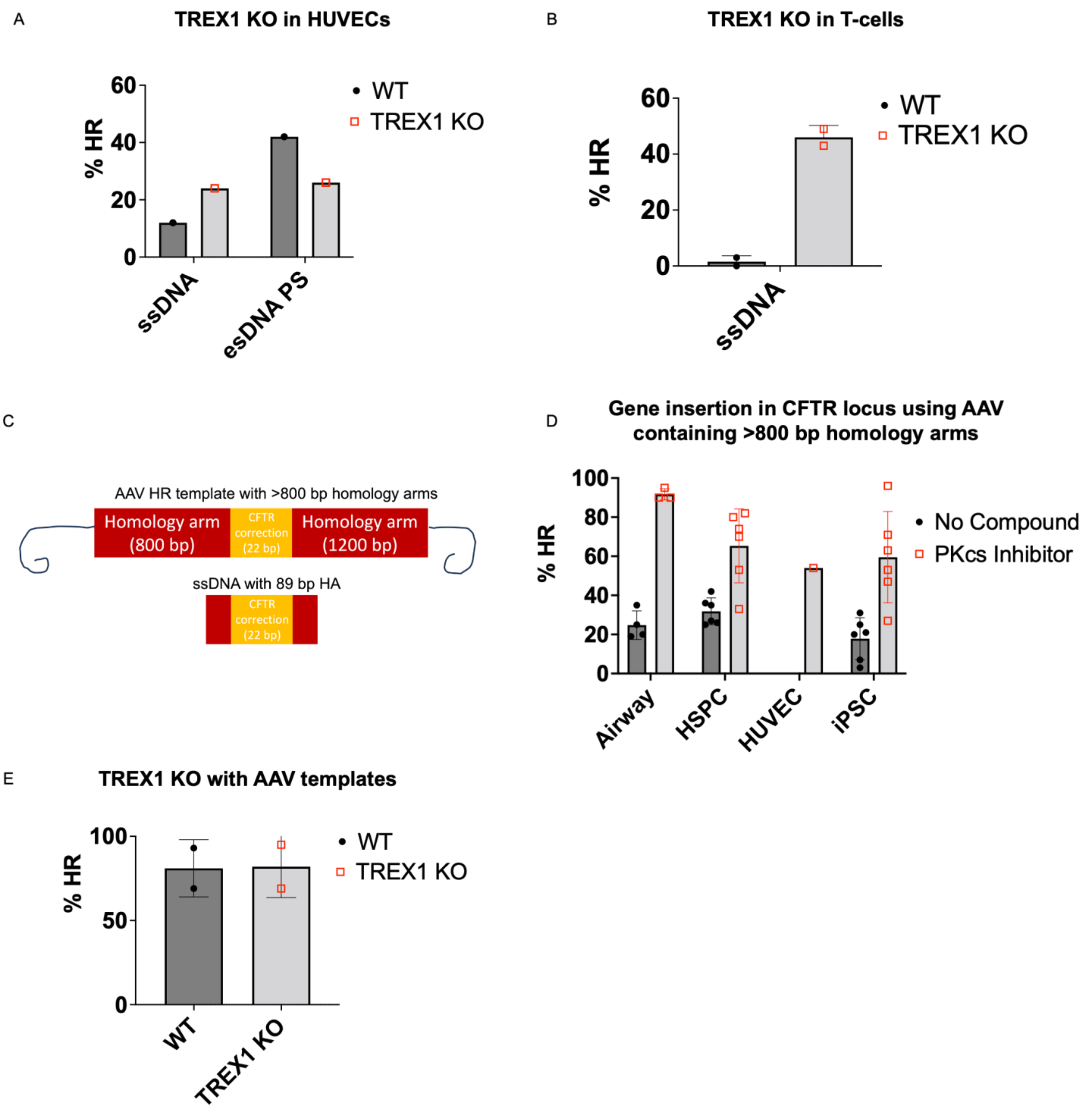


**Figure S3**: Gene insertion using AAV and Cas9 Targeting Sequences

**A**) Knocking out *TREX1* in HUVECs improved gene insertion using unmodified ssDNA from 12% to 24% alleles with HR in the *CFTR* locus (n = 1 biological replicate). **B**) Knocking out *TREX1* in T-cells improved gene insertion using unmodified ssDNA from 1.5% to 46% alleles with HR in the *CFTR* locus (n = 1 biological replicate). **C**) Graphic comparing AAV-based HR templates carrying the F508del correction sequence with long homology arms and ssDNA templates with short homology arms. Both reagents target exon 11 of the *CFTR* locus. **D**) Gene insertion using AAV templates was efficient in all cell types tested. The addition of DNA-PKcs inhibitor improved gene insertion in all cell types. We did not edit HUVECs in the absence of DNA-PKcs inhibition. **E**) Knocking out *TREX1* did not affect gene insertion using AAV-based templates targeting the *CFTR* locus.
